# Retrotransposon insertions can initiate colorectal cancer and are associated with poor survival

**DOI:** 10.1101/443580

**Authors:** Tatiana Cajuso, Päivi Sulo, Tomas Tanskanen, Riku Katainen, Aurora Taira, Ulrika A. Hänninen, Johanna Kondelin, Linda Forsström, Niko Välimäki, Mervi Aavikko, Eevi Kaasinen, Ari Ristimäki, Selja Koskensalo, Anna Lepistö, Laura Renkonen-Sinisalo, Toni Seppälä, Teijo Kuopio, Jan Böhm, Jukka-Pekka Mecklin, Outi Kilpivaara, Esa Pitkänen, Kimmo Palin, Lauri A. Aaltonen

## Abstract

Genomic instability pathways in colorectal cancer (CRC) have been extensively studied, but the role of retrotransposition in colorectal carcinogenesis remains poorly understood. Although retrotransposons are usually repressed, they become active in several human cancers, in particular those of the gastrointestinal tract. Here we characterize retrotransposon insertions in 202 colorectal tumor whole genomes and investigate their associations with molecular and clinical characteristics. We found highly variable retrotransposon activity among tumors and identified recurrent insertions in 15 known cancer genes. In approximately 1% of the cases we identified insertions in *APC*, likely to be tumor-initiating events. Insertions were positively associated with the CpG island methylator phenotype and the genomic fraction of allelic imbalance. Clinically, high number of insertions was independently associated with poor disease-specific survival.

## INTRODUCTION

Retrotransposons are transposable genetic sequences that copy themselves into an RNA intermediate and insert elsewhere in the genome. Almost half of the human genome consists of transposon derived sequences ^1^, however only a few elements remain retrotransposition competent and account for most retrotranspositions ^2,3^. Two types of retrotransposons have been identified in the human genome; autonomous and non-autonomous. Autonomous elements, such as Long Interspersed Nuclear Element-1s (LINE-1s) and Endogenous retroviruses (ERVs), provide the required machinery for retrotransposition. On the contrary, non-autonomous elements, such as Alus and SINE-VNTR-Alu (SVAs), require the LINE-1 machinery to retrotranspose ^4–7^. In cancer, approximately 24% of somatic retrotranspositions involve 3′transduction, a process characterized by mobilization of 3′ flanking sequence which can serve as a unique sequence revealing the insertion origin ^8–11^.

LINE-1s are frequently repressed by promoter methylation ^12^ and genome-wide hypomethylation is reported to lead to their activation during tumorigenesis ^13,14^, thus leading to high retrotransposon activity and genome instability ^15,16^. High retrotransposon activity has been reported in several human cancers, especially in tumors arising from the gastrointestinal tract, such as colorectal cancer (CRC) ^10,11,17–20^. Somatic insertion density in tumors is higher in closed chromatin and late replicating regions. Among insertions in genes, insertion density is higher in genes with low expression ^10,21^. Furthermore, ongoing retrotransposon activity has been reported in CRC ^22^. Insertion count is associated with patient age ^18^ and LINE-1 hypomethylation is associated with poor survival in colorectal cancer ^23^. LINE-1 insertions in *APC* have been reported in two CRCs, indicating that these insertions may be early tumorigenic events ^24,25^.

CRC can develop through two distinct pathways; chromosomal instability (CIN) or microsatellite instability (MSI). Most sporadic CRCs follow the CIN pathway, characterized by a large number of chromosomal alterations. Fifteen percent of CRC cases follow the MSI pathway, characterized by a high number of base substitutions and short insertions and deletions ^26^. Seventy-five percent of MSI-positive sporadic CRCs are attributed to the CpG island methylator phenotype (CIMP) ^27^ which is characterized by gene promoter hypermethylation. Although genomic instability pathways have been studied extensively in CRC, the tumorigenic role of retrotransposition is not fully understood. Retrotransposon insertions have been difficult to detect with previous methodological approaches and very few genome-wide studies have been reported. Here, we characterized somatic retrotransposon insertions in 201 colorectal cancers and one colorectal adenoma utilizing whole genome sequencing (WGS), and investigated the associations between somatic retrotransposon activity and clinical characteristics.

## RESULTS

### Genome-wide detection of somatic retrotransposition in colorectal tumors

To characterize the landscape of somatic retrotransposon insertions in CRC we applied TraFiC ^10^ and DELLY ^28^ to WGS data from 202 colorectal tumors and matched normal samples. From the 202 tumors, 12 were MSI and 190 were microsatellite stable (MSS) including 3 ultra-mutated tumors, harboring somatic *POLE* mutations. After strict somatic filtering, we identified a total of 5,072 insertions **(Supplementary Table S1)**. Based on visual inspection of the paired-end read data on 100 random insertion calls, 76 calls were evaluated as true somatic insertions, giving a false positive rate of 24% (95% confidence interval [CI], 16-34%) (**Supplementary Table S1)**. Additionally, 14 out of 15 3`transductions from two samples were validated by long-distance inverse-PCR (LDI-PCR) and Nanopore sequencing in a separate study ^22^. The mean number of insertions per tumor was 25 (median, 17; interquartile range, 10-31) with high variability among tumors (Figure 1a). Mean number of insertions in MSS, MSI and the *POLE* ultra-mutated tumors was 25, 34 and 24 respectively. The majority of insertions (99%, 5024/5072) were LINE-1 retrotranspositions, however we also detected 20 SVA, 13 Alu and 15 ERV insertions (**Supplementary Table S1**). In concordance with previous studies ^10,21^, insertion density was higher in closed chromatin (1.78 insertions per Mbp) than in open chromatin (0.96 insertions per Mbp) and in late replicating regions (replication time>0.8, 3.06 insertions per Mbp) than in early replicating regions (replication time<0.2, 0.73 insertions per Mbp) (Figure 1b).

**Figure 1.**
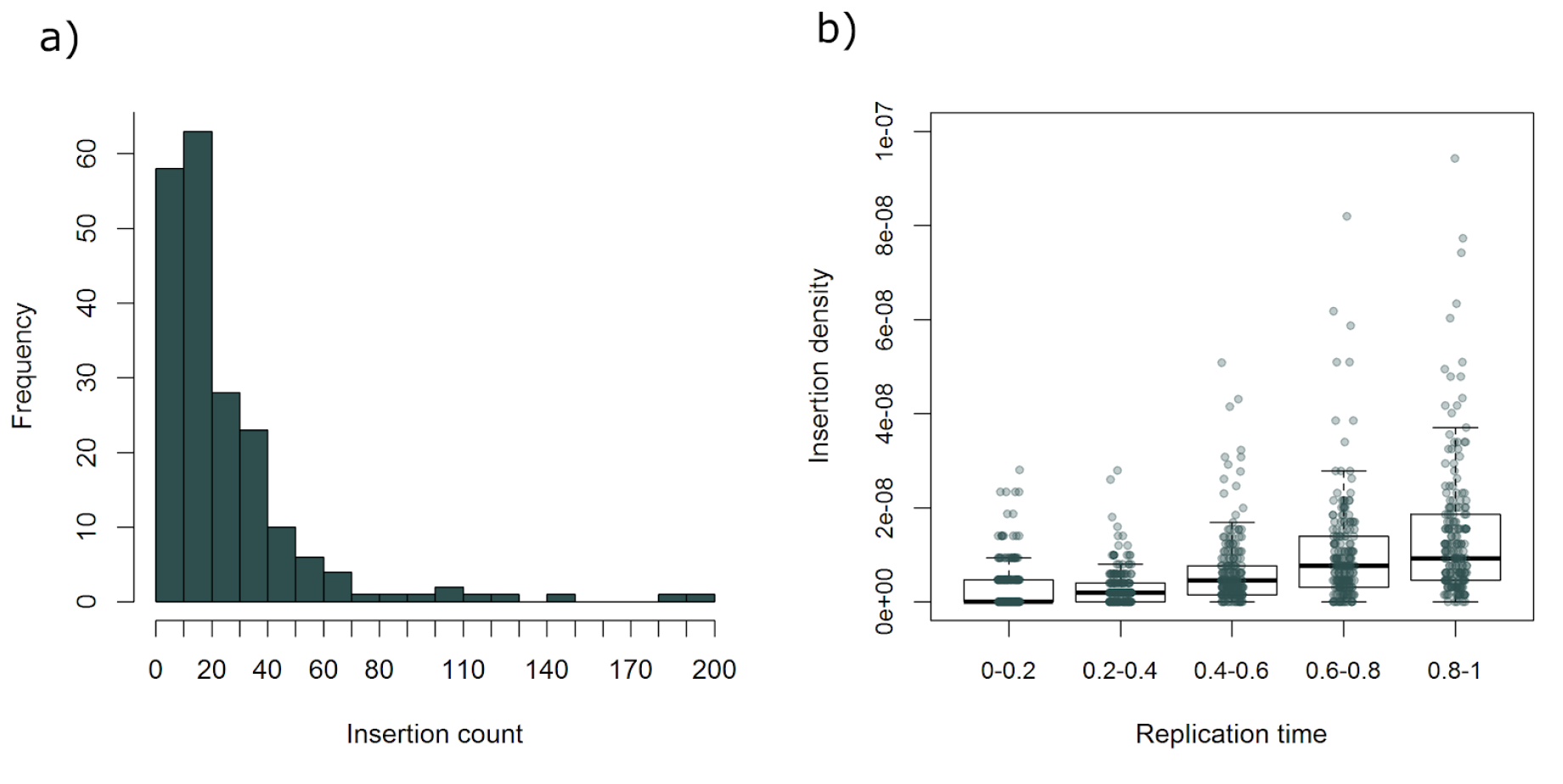
Distribution of somatic insertions across 202 colorectal tumors and over replication time. a) Frequency of somatic insertion counts in 202 colorectal tumors. b) Insertion density over replication time. The genome was stratified by replication time in five categories where 0 referred to the earliest replication timing. Each point represents insertion density in the corresponding category for each of the 202 tumors.

### Retrotransposon insertions are initiating events in approximately 1% of colorectal tumors

To characterize retrotransposon insertions in genes, all protein-coding transcripts and the insertion polyA/T in conjunction with gene orientation were used to assess insertion orientation. Of the 5,072 insertions, 1,680 (33%) were detected within protein-coding genes, with 98% in introns (**Supplementary Table S1, Supplementary Figure S1**). We identified 399 insertions in sense orientation and 472 in antisense orientation, suggesting antisense preference (Exact Binomial Test 95% CI, 50.8-57.3%, p=0.014) (**Supplementary Table S1**). Insertion count was higher in genes with lower expression (median transcript per million reads [TPM] from 34 tumors) in concordance with a previous study ^21^ (Figure 2a). Recurrent insertions (at least two insertions) were identified in 333 protein-coding genes (**Supplementary Table S2**). The most enriched biological processes among these genes were neuron-neuron synaptic transmission and cell-cell adhesion (Figure 2b).

**Figure 2.**
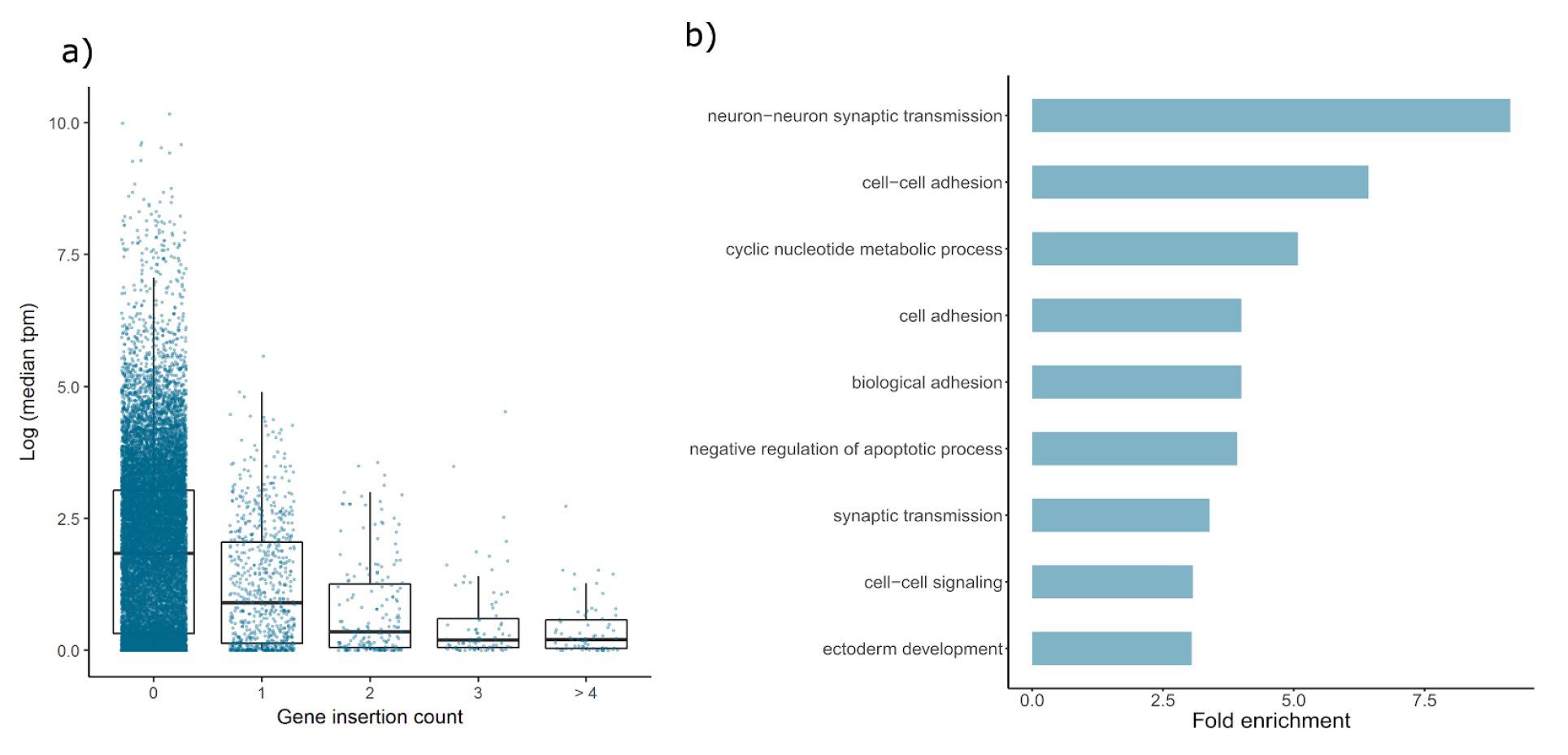
Retrotransposon insertions in protein-coding genes. a) Gene expression (median TPM values from 34 tumors) over gene insertion count groups. b) Biological processes significantly enriched at least 2 fold in protein-coding genes with two or more insertions.

Fifteen genes in the Cancer Gene Census (CGC) ^29^ displayed recurrent insertions (Table 1). The most frequently affected protein-coding gene was *LRP1B* with 19 insertions, all located in introns. *LRP1B* is a known fragile site ^30^ and has been classified as a tumor suppressor gene in the CGC. No clear bias towards tumor suppressor genes or oncogenes was apparent among the recurrent targets (Table 1). We also investigated whether insertions had an overall effect on the expression of the closest genes but no significant effect was detected (**Supplementary Figure S2, Methods section “Association test between insertions and RNA expression”**).

**Table 1.**
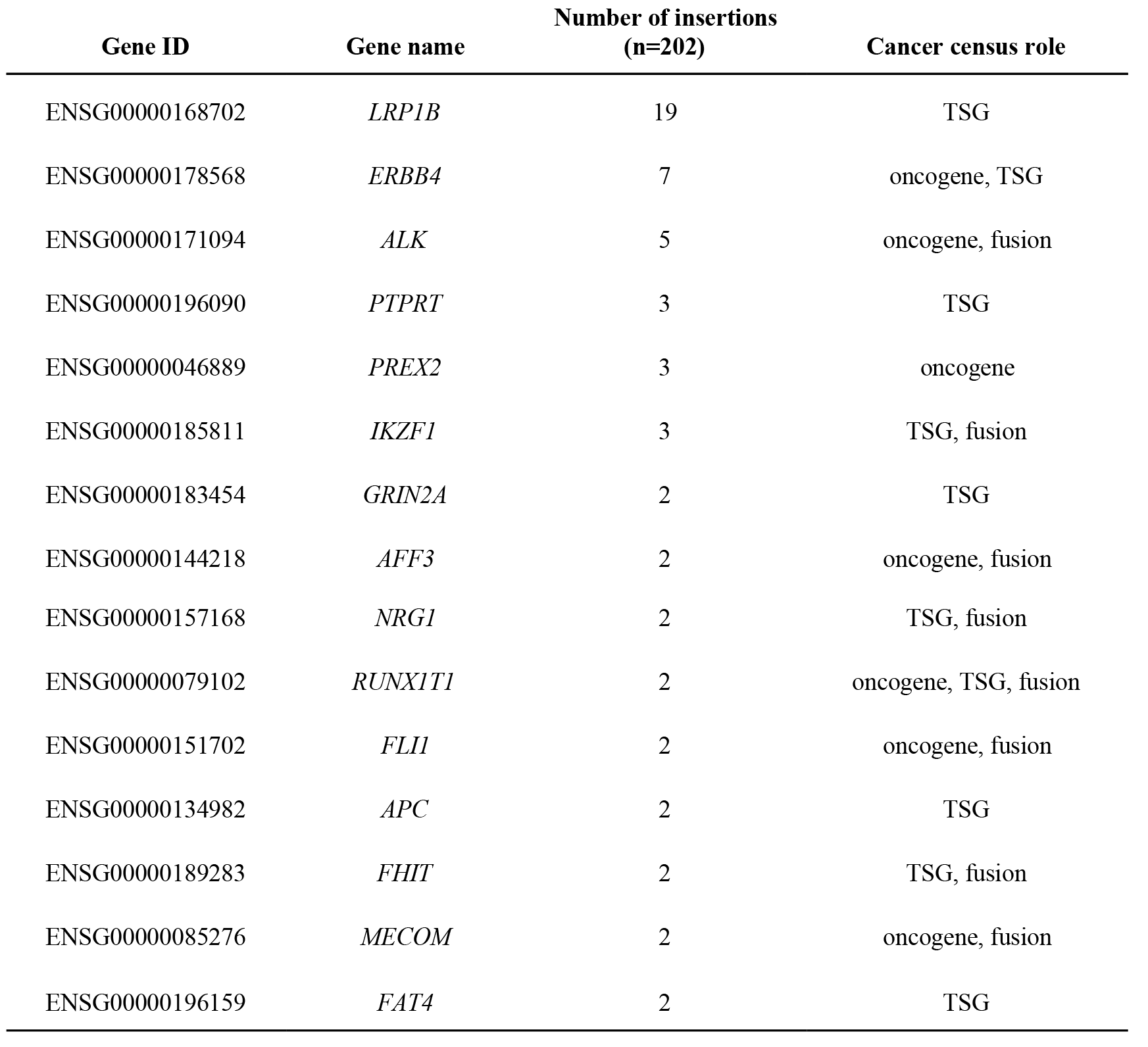
Genes from the Cancer Gene Census with two or more insertions; Cancer census role, role in cancer as defined by the Cancer Gene Census ^29^. TSG, tumor suppressor gene.

Seventy-two insertions were identified in the exons of protein-coding genes (**Supplementary Table S1**, **Supplementary Figure S1**). We identified one insertion in the last exon/3′UTR of *PIK3CA* (**Supplementary Table S1**) and two insertions in exon 16 of *APC* (Figure 3**, Supplementary Table S1**). Loss of heterozygosity and copy number loss encompassing *APC* were found in both tumors, and no other sequence variations were identified. Moreover, both insertions were in close proximity (2,151 bp) to two previously reported insertions ^31,32^ and the location of the insertions was consistent with the distribution of non-synonymous point mutations detected in *APC* (Figure 3). Altogether, these findings suggest that retrotransposon insertions contributed to the early steps of tumorigenesis in 2 of the 202 colorectal tumor patients.

**Figure 3.**
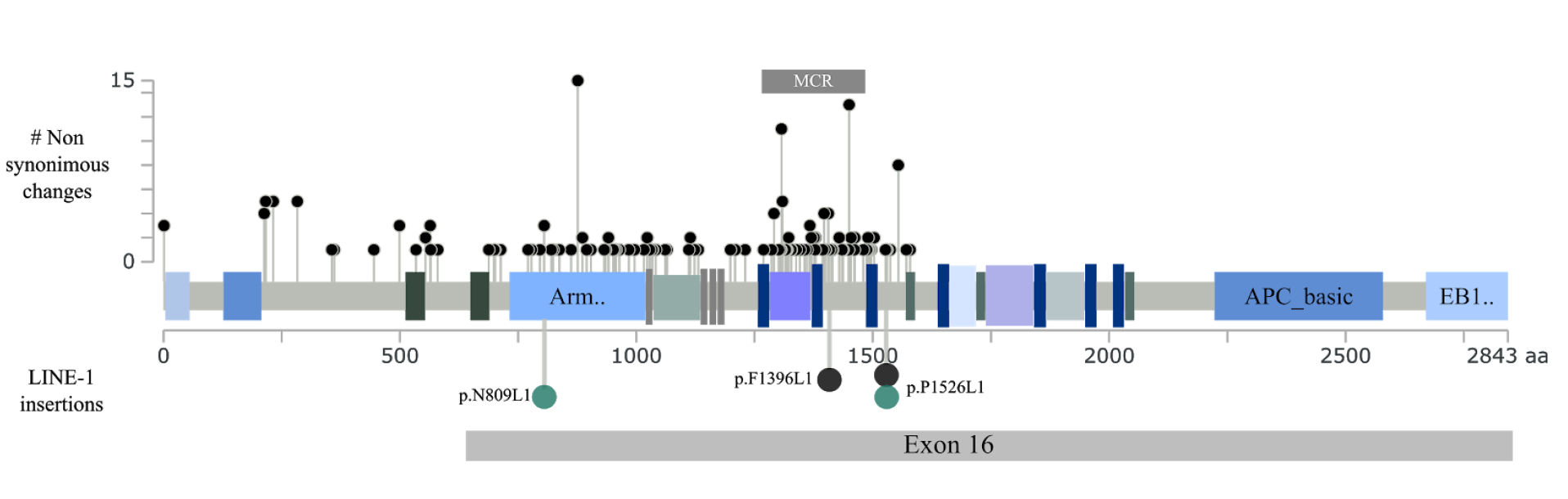
Distribution of non-synonymous changes and LINE-1 insertions on the linear protein of *APC*. Non-synonymous changes in 187 MSS CRCs, small lollipops. LINE-1 insertions, larger lollipops. p.N809L1 (c1049-1T) and p.P1526L1 (c310.1T), turquoise lollipops; p.F1396L1 and p.P1526L1 ^31,32^, black lollipops. Figure modified from cBio cancer genomics portal ^33,34^.

### Recurrent insertions in lowly expressed chromosomal fragile sites

We observed recurrent insertions in 12 out of 21 fragile sites ^30^ (**Supplementary Table S3**). Since common fragile sites are prone to copy number alterations (CNAs)^35^, we evaluated whether retrotransposition and CNAs - in this study detected as allelic imbalances (AI) – were correlated (**Supplementary Table S3**). Fragile sites with high frequency of insertions seemed to display lower frequency of allelic imbalances (Figure 4a). Next, we investigated whether this difference could result from differences in gene expression within fragile sites. Indeed, insertion frequency seemed to be higher in genes with lower expression (Exact Two-Sample Fisher-Pitman Permutation Test for log-transformed gene expression, p=0.03863) (Figure 4b). These results are concordant with our data and those of another study ^21^; insertion density is overall negatively correlated with gene expression.

**Figure 4.**
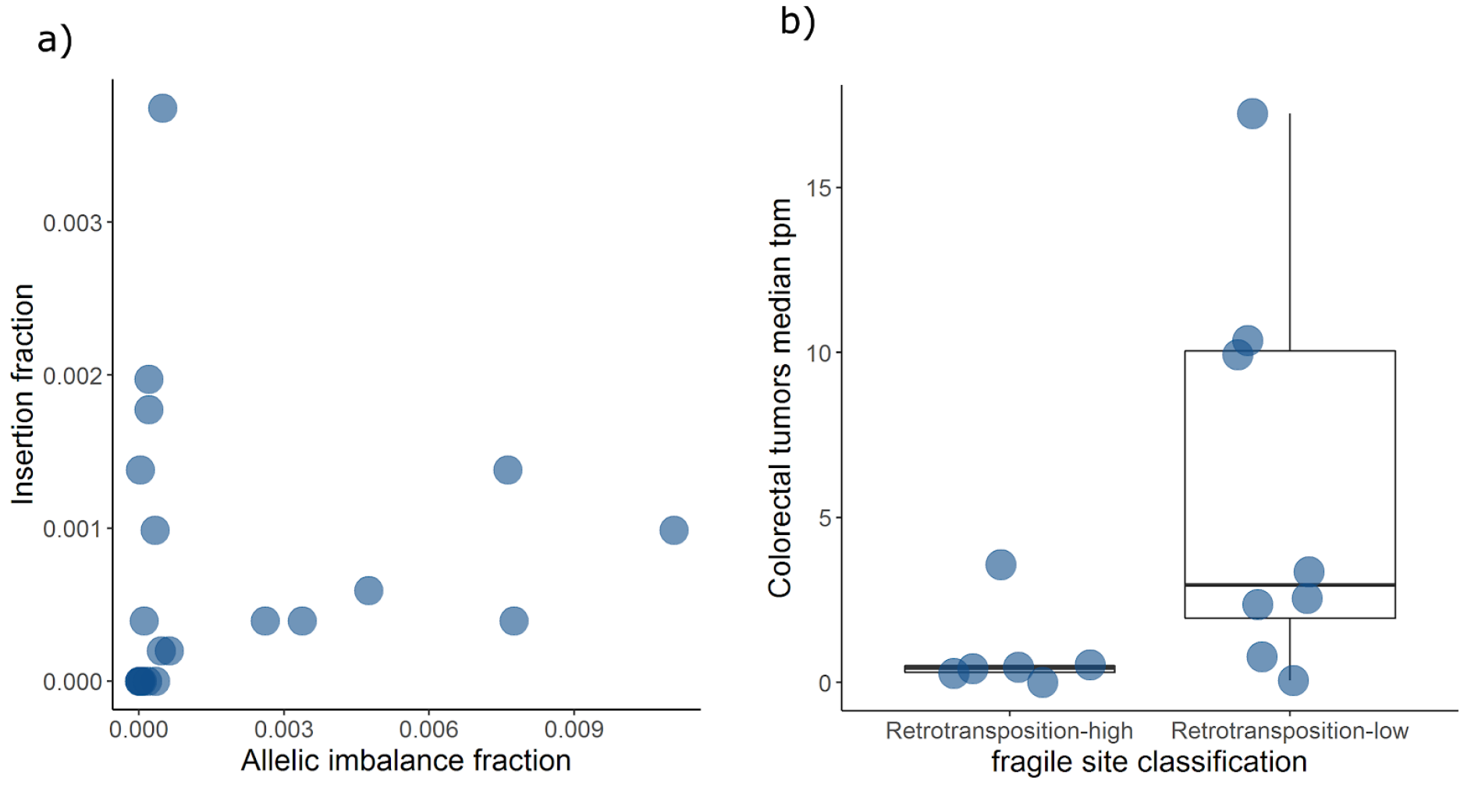
Insertion and AI frequency in 21 fragile sites. a) Insertion fraction over the fraction of allelic imbalance in 21 fragile sites. b) Gene expression (median TPM values from 34 tumors) over fragile site classification based on the ratio of insertion fraction and fraction of allelic imbalance events (**Supplementary Table S3**).

### Few active LINE-1s account for the majority of LINE-1 transductions

We utilized the 3′unique sequence from the transduced regions to identify the source elements of LINE-1 transductions. We detected a total of 347 transductions arising from 56 of 315 human specific full-length LINE-1s. Recurrent transductions were detected from 24 LINE-1s, and in concordance with our previous study ^11^ the most active was the LINE-1 located in 22q12.1, which alone accounted for 160 transductions (46%). Seven and six percent of the transductions arose from the LINE-1s located in 9q32 and Xp22.2, respectively. The active LINE-1s reported in this study are in concordance with the frequencies reported by another study across 31 different tumor subtypes (**Supplementary Table S4**) ^21^.

### Insertion count is significantly associated with CIMP and the genomic fraction of allelic imbalance

We investigated the associations between insertion counts and molecular and clinical characteristics. We utilized 196 colorectal tumors with complete information on molecular and clinical variables that were included in the model (Table 2, **Supplementary Table S5**). We applied a multiple linear regression model for log-transformed insertion counts, and hypothesized that the number of somatic insertions may be associated with tumor location, *TP53* mutation, MSI, genomic fraction of allelic imbalance and CIMP. The model was adjusted for mean sequencing coverage, tumor stage, sex and age at diagnosis (Table 2). Goodness-of-fit was tested by Pearson’s chi-square test (p=0.99). We found that insertion count was positively associated with CIMP (p=0.00032) and the genomic fraction of allelic imbalance (p=0.0036) (Table 2).

**Table 2.** Multiple linear regression model for log insertion counts. MSI, microsatellite instability; CIMP-H, CpG methylator phenotype high. Significance codes: 0.0001‘***’ 0.001‘**’ 0,01‘*’ 0,05‘.’

|  | Coefficient | Std. Err. | z | p | Signif. |
| --- | --- | --- | --- | --- | --- |
| Intercept | 0.408 | 0.647 | 0.630 | 5.29e-01 |  |
| CIMP-H | 0.607 | 0.169 | 3.60 | 3.22e-04 | *** |
| Allelic Imbalance (/10% of reference) | 0.0826 | 0.0284 | 2.91 | 3.64e-03 | ** |
| *TP53* mutation | -0.0684 | 0.134 | -0.509 | 6.10e-01 |  |
| MSI | 0.150 | 0.285 | 0.527 | 5.98e-01 |  |
| Mean coverage (/10 reads) | 0.309 | 0.0862 | 3.59 | 3.33e-04 | *** |
| Age at diagnosis (/10 years) | 0.0331 | 0.0603 | 0.549 | 5.83e-01 |  |
| Male | 0.0570 | 0.123 | 0.464 | 6.42e-01 |  |
| Dukes B | -0.0578 | 0.168 | -0.344 | 7.31e-01 |  |
| Dukes C | -0.0483 | 0.186 | -0.259 | 7.96e-01 |  |
| Dukes D | 0.00490 | 0.211 | 0.0232 | 9.81e-01 |  |
| Proximal location | 0.206 | 0.144 | 1.43 | 1.52e-01 |  |

### Insertion count is significantly associated with poor disease-specific survival

We applied the Cox proportional hazards model in 192 patients with complete information on molecular and clinical variables that were used in the model (Table 3**, Supplementary Table S5**). Patients were followed for 1,370 person-years (**Supplementary Table S5**). We hypothesized that insertion counts may be associated with disease-specific survival (Figure 5). The model was adjusted for tumor stage, sex, MSI, the genomic fraction of allelic imbalance, *BRAF* mutation and CIMP status (Table 3). As expected, advanced tumor stage (Dukes C and D) was strongly associated with CRC-specific survival. However, even after adjusting for the above-mentioned covariables, insertion count was independently associated with poor disease-specific survival (p=0.0029) (Figure 5, Table 3).

**Figure 5.**
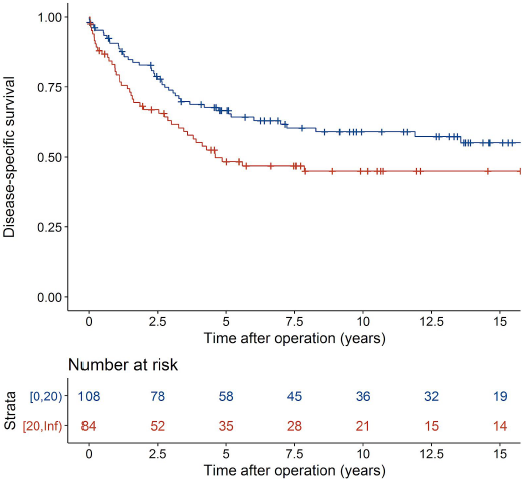
Kaplan-Meier curves by insertion count. Tumor with less than 20 somatic insertions (blue line) and tumor with 20 or more insertions (red line).

**Table 3.** Cox proportional hazards model for disease-specific survival. The model was stratified by tumor location. HR, Hazard ratio; CI, confidence interval; MSI, microsatellite instability; CIMP-H, CpG island methylator phenotype high. Significance codes: 0.0001‘***’ 0.001‘**’ 0,01‘*’ 0,05‘.’

|  | Coefficient | Std. Err. | z | p | HR [95% CI] | Signif. |
| --- | --- | --- | --- | --- | --- | --- |
| Insertion count (/10) | 0.108 | 0.0362 | 2.98 | 2.93e-03 | 1.11 [1.04, 1.20] | ** |
| MSI | -0.258 | 0.642 | -0.402 | 6.88e-01 | 0.773 [0.219, 2.72] |  |
| CIMP-H | 0.174 | 0.341 | 0.510 | 6.10e-01 | 1.19 [0.610, 2.32] |  |
| *BRAF* mutation | 0.790 | 0.447 | 1.77 | 7.68e-02 | 2.20 [0.918, 5.29] | . |
| Age [55, 75) years | -0.147 | 0.408 | -0.360 | 7.19e-01 | 0.863 [0.388, 1.92] |  |
| Age ≥ 75 years | 0.188 | 0.427 | 0.439 | 6.60e-01 | 1.21 [0.523, 2.78] |  |
| Male | 0.311 | 0.232 | 1.34 | 1.80e-01 | 1.37 [0.866, 2.15] |  |
| Dukes B | 0.452 | 0.449 | 1.01 | 3.13e-01 | 1.57 [0.652, 3.79] |  |
| Dukes C | 1.77 | 0.431 | 4.12 | 3.82e-05 | 5.89 [2.53, 13.7] | *** |
| Dukes D | 2.78 | 0.454 | 6.12 | 9.07e-10 | 16.2 [6.64, 39.4] | *** |
| Allelic Imbalance (/10% of reference) | -0.0583 | 0.0539 | -1.08 | 2.80e-01 | 0.943 [0.849, 1.05] |  |

## DISCUSSION

Although retrotransposon activity is a hallmark of tumors of the gastrointestinal tract ^10,11,17–20^, the role of retrotransposon insertions in CRC remains unclear with very few studies reported. Here, we characterized the somatic landscape of retrotransposon insertions in the largest dataset of colorectal tumor whole-genomes reported to date, and identified significant associations with clinical characteristics.

We observed high retrotransposon activity with wide variability among tumors. We confirmed higher insertion density in late replicating regions, closed chromatin. Among insertions in genes, we also observed higher insertion count in genes with lower expression. The list of the most active LINE-1s became also validated in this extended set of CRCs ^10,11,21^. A number of novel observations were made. We identified recurrent insertions in 333 protein-coding genes, 15 of which are included in the Cancer Gene Census ^29^. The most recurrent hit was *LRP1B* with 19 intronic insertions. The high frequency of insertions in this gene could be a result of various characteristics such as chromatin state, replication timing as well as gene length and/or expression. However, other causes such as sequence composition or somatic selection cannot be excluded. We also observed a high frequency of insertions at fragile sites with lower gene expression and lower allelic imbalance fraction. These findings suggest that while allelic imbalance is a recurrent feature of some fragile sites, sites with lower gene expression are more prone to retrotransposon insertions in CRC.

Among the exonic insertions, we identified one in *PIK3CA* and two in *APC*. *PIK3CA* is a known oncogene involved in colorectal tumor progression and mutations in *APC* lead to colorectal tumor initiation ^36^. The insertions in *APC* were consistent with the distribution of non-synonymous changes in this gene and two previously reported insertions ^31,32^, indicating that retrotransposon insertion in *APC* is one mechanism of CRC initiation.

The availability of patient data allowed us to investigate possible associations between somatic insertion count and various molecular and clinical characteristics. We applied a multiple linear regression model and found that retrotransposon activity was positively associated with the genomic fraction of allelic imbalance, and paradoxically with CIMP even though LINE-1s are frequently repressed by promoter methylation ^12^. Of note, both CIMP and the genomic fraction of allelic imbalance are characteristic of the two distinct genetic instability pathways in CRC. No associations with age at diagnosis, *TP53* mutations or tumor stage were detected in contrast to other studies ^18,37^. These studies had significantly smaller sample sizes and fewer covariables were taken into account, which may explain the discrepancy. Importantly, survival analysis revealed a significant association between insertion count and poor disease-specific survival independently of other prognostic factors. Our findings indicate that tumors with high retrotransposon activity present characteristics of both MSS and MSI tumors, and are associated with poor CRC-specific survival.

By characterizing the landscape of retrotransposon insertions in a large dataset of CRCs, we identified that retrotranspositions can serve as tumor initiating events in CRC. The association of retrotransposition events with clinical characteristics - in particular poor prognosis - suggest that retrotransposition may play a more important role in CRC than previously thought. Further work should elucidate the timing and mechanisms leading to high somatic retrotransposition activity in some individuals, while others are spared. Understanding these could provide new tools for management of CRC, including prevention.

## METHODS

### Study subjects

The samples and the clinical data utilized in this study were obtained from a population based series of 1042 CRCs described previously ^38,39^ and from a subsequently collected series of 472 additional Finnish CRCs. The tumors were fresh frozen and the corresponding normal tissues were obtained from either blood or from the normal colon tissue. Originally 202 CRCs entered the analyses. However, one tumor was later classified as an advanced adenomatous lesion (c232.1T). The study was reviewed and approved by the Ethics committee of the Hospital district of Helsinki and Uusimaa, Finland. Either a signed informed consent or authorization from the National Supervisory Authority for Welfare and Health was obtained for all the samples.

### Whole genome sequencing

Whole genome sequencing was performed on Illumina HiSeq 2000 with 100 bp paired-end reads. Each normal and tumor DNA was sequenced to at least 40x median coverage as described in previous studies ^40^ (Palin K et al., Nature Communications, 2018, in press).

### Transposon detection

The identification of somatic retrotransposon insertions was conducted utilizing the Transposon Finder in Cancer (TraFiC) ^10^. TraFiC default parameters were applied except for; a=1 (RepeatMasker accuracy), s=3 (minimum of 3 reads in tumor cluster), and gm=3 (minimum of 3 reads in normal cluster). In addition, paired-end reads with both ends having equal mapping quality and above 0 were included. In these cases, the first end of the pair was selected as the anchor read (end mapping to non-repetitive sequence). RepeatMasker (version open-4.0.5) and NCBI/RMBLAST 2.2.27+ were used for retrotransposon alignment as part of TraFiC. The RepeatMasker Database release utilized in this study was 20140121 ^41,42^. Somatic filtering was performed against germline calls from 234 normal samples (202 corresponding CRC normals, 20 myometrium samples ^43^ and 12 blood samples ^44^) with a 200 bp window as described in TraFiC ^10^. Furthermore, calls in decoy sequences from 1000 Genomes Project Phase 2 (hs37d5) ^45^ were filtered away.

### Detection of LINE-1 transductions

We identified 3’ and orphan transductions utilizing DELLY structural variant (SV) calls (v 0.0.9) ^28^ from a previous study ^40^. Filtering criteria utilized in this study were: SV calls supported by at least three supporting discordant reads, mapping quality >37, SV length >1000 bp. Subsequently, we extracted the SVs with one end of the pair within 1000 bp from the 3’ end of a reference human-specific LINE-1 (Reference L1HS, full-length) from The European database of L1-HS retrotransposon insertions in humans (euL1db) ^46^, database version v1.0, date 05-10-14. The other end of the pair was used in the somatic filtering, where a 200 bp window and transduction calls from the pool of normal samples above mentioned were applied. One transductions detected in a female coming from an LINE-1 in Yp11.2 was filtered away. Furthermore, transduction calls within 200 bp, from the same retrotransposon family, and in the same sample were regarded as the same insertion and merged together. The same rationale was applied for calls detected by both DELLY and TraFiC.

### Allelic imbalance

The allelic imbalance calls derived from a separate study (Palin K et al., Nature Communications, 2018, in press). In brief, both the tumor and respective normal DNA were genotyped with Infinium Omni2.5-8 (Illumina Inc.) array at the Estonian Genome Center. The B-Allele Frequencies and Log-R ratios were extracted with Illumina GenomeStudio software and the allelic imbalance regions were calculated for all samples using BAF segmentation ^47^ with default parameters.

### Methylation-Specific Multiplex Ligation-dependent Probe Amplification (MS-MLPA)

An MS-MLPA assay (Nygren AO, 2005) with the SALSA MLPA ME042 CIMP probemix (MRC-Holland, Amsterdam, The Netherlands) was used to determine the CpG island methylator phenotype (CIMP) in an extended set of 255 tumor samples and the corresponding normal colon tissue of 175 samples as a separate study. Data from normal samples were used to determine the threshold for hypermethylation in the tumor samples. MS-MLPA was performed according to manufacturer’s instructions 48 (http://www.mrc-holland.com Accessed December 2015). In short, the assay targets the promoter region of 8 tumor suppressor genes; *CACNA1G*, *CDKN2A*, *CRABP1*, *IGF2*, *MLH1*, *NEUROG1*, *RUNX3*, *SOCS1*. The methylation level for each probe was called using the Coffalyser software (MRC-Holland, Amsterdam, The Netherlands). If ≥25% of the probes for one gene were methylated, the gene was scored as methylated. If 5-8 genes were scored as methylated, the tumor was classified as a CIMP-high (CIMP-H), and if 0-4 genes were scored as methylated it was classified as a CIMP-low (CIMP-L) tumor.

### RNA sequencing

Total RNA from consecutive cryosections was extracted using RNeasy Mini Kit (Qiagen) from 34 tumors that displayed more than 50% of cancer cell percentage (HE staining of cryosections) and RNA integrity>6 (Agilent RNA 6000, Agilent 2100 Bioanalyzer). RNA-seq data was processed using Kallisto (version 0.43.0) software ^49^. Kallisto quantification was executed in paired-end mode and aligned against the Ensembl Human reference transcriptome (GRCh37_79). Quantification results from Kallisto were normalized and aggregated to gene-level utilizing sleuth (version 0.28.1) R package ^50^ with default filtering settings ^50^.

### Visual inspection of paired-end read data

We selected 100 random insertions to ascertain the rate of true somatic calls based on visual inspection of the paired-end read data. Visualization was performed with BasePlayer ^51^. Somatic calls were visually validated as true if the insertion call was supported by discordant reads (three+three for TraFiC calls) and at least two split reads supporting the insertion breakpoint and/or the polyA/T. Furthermore, the corresponding normal tissue was also visualized to confirm the somatic origin of the insertion calls.

### Insertion annotation

Annotation of the insertion calls was applied by using the inner genomic coordinates of the reciprocal clusters provided by TraFiC (P_R_POS & N_L_POS) (**Supplementary Table S1**). Insertion breakpoints hitting an intron or an exon of any protein-coding transcript (GRCh37_87) were annotated as intron/exon hit. Insertion orientation was determined by the presence of a polyA or a polyT in conjunction with gene orientation. To call a polyA, at least two forward split reads with at least five consecutive As at the 5′end of the read were required. To call a polyT, at least two reverse split reads with at least 5 consecutive Ts at the 3′end of the read were required. Sense insertions were defined when insertion orientation and gene orientation were the same whereas antisense insertions were defined when insertion and gene orientation were the opposite. Replication time fractions were extracted from another study ^52^. Insertion density was defined as number of insertions divided by the total number of base pairs of each replication time fraction. Open chromatin was defined as DNAse regions that were overlapping in at least two out of the four cell lines (RKO, LoVo, CaCo2 and Gp5D) (GSE83968) ^53^ in the 1000 Genomes Project PilotMask reference genome ^54^. Closed chromatin regions were defined as the above-mentioned reference genome minus the open chromatin regions.

### Gene ontology analysis

Gene ontology enrichment analysis was performed with PANTHER Overrepresentation Test (Released 20171205), annotation version 13.1 released 2018-02-03 and the test utilized was Fisher’s exact test with FDR multiple test correction ^55^.

### Fragile sites

The 21 fragile sites were defined as genes with more than 0.85 probability of being fragile (Random forest 3 predictors) as estimated in another study ^30^. Genomic coordinates were lifted to GRCh37/Hg19 with https://genome.ucsc.edu/cgi-bin/hgLiftOver and regions with no converted coordinates were excluded (chr10:46597226-48877831, chr10:45970128-48447930). The fraction of AI was calculated as the number of focal AI events *per* fragile site (both breakpoints of each AI call within the fragile site coordinates) divided by the total number of AI events in 1699 tumors (Palin K et al., Nature Communications, 2018, in press). Insertion fraction was calculated as the number of insertions per fragile site divided by the total number of insertions detected in 202 patients. Fragile site categories were defined based on the ratio of insertions/AI. Retrotransposon-low; 0 < ratio < 1, and Retrotransposon-high; ratio > 1.

### Mutation analysis in *BRAF*, *KRAS*, *TP53* and *APC*

Somatic changes in *BRAF, KRAS, TP53* and *APC* were called using MuTect (version 1.1.4) with default parameters (GRCh37_78), as described in a previous study ^40^ (Palin K et al., Nature Communications, 2018, in press). Subsequent filtering criteria were minimum coverage of 4, minimal allelic fraction of 10 and minimum quality score 20 ^51^. For *KRAS*, mutations in codons 12, 13, 61, 117 and 146 in any transcript were classified as mutation positive and for *BRAF*, only hotspots in V600E in any transcript were considered as mutation positive. All non-synonymous changes in any transcript of *TP53* were classified as mutation positive. In addition, non-synonymous changes in *APC* (ENST00000457016) from 234 MSS tumors (Palin K et al., Nature Communications, 2018, in press) were utilized for Figure 3. Figure 3 was created with http://www.cbioportal.org/tools.jsp ^33,34^ and modified with Inkscape (http://www.inkscape.org).

### Association test between insertions and RNA expression

For the 827 insertions identified in any of the 34 tumors, we investigated the effect on the expression of the 642 distinct closest genes. For each sample and each gene the TPM values were extracted and ranked in an ascending order. Consequently, the rank number corresponding to the sample with the insertion was recorded for each gene. We computed the sum-of-squared error statistic (Chi-square test) for the frequency table to test whether the rank values of the samples with insertion were uniformly distributed (no insertion effect on gene expression) (**Supplementary Figure S2**). Furthermore, 100 000 permutations with randomized rank numbers were applied but no significant effect was observed (**Supplementary Figure S2**). Tests were performed using R versions 3.4.3 or 3.3.0.

### Multiple linear regression analysis

To model retrotransposon insertion counts we applied a multiple linear regression model for log-transformed insertion counts. Spearman correlation matrix (R package PerformanceAnalytics) and variance inflation factors (vif function in R package car) were computed to evaluate possible collinearity among explanatory variables ^56,57^. Model fit was assessed by plotting residuals against fitted values, theoretical normal quantiles and leverage (**Supplementary Figures S3-S5**). All tests were performed using R version 3.3.2 ^58^.

### Cox proportional hazards regression analysis

We applied the Cox proportional hazards regression to study the association between disease-specific survival with retrotransposon insertion counts. The time variable was defined as days since diagnosis or operation. Patients that were alive in the last status assessment were censored at that date (survival status was assessed periodically using the Population Register Centre of Finland with the most recent assessment in 2016). Death from other causes than CRC were also defined as censored events. Proportional hazards assumptions were assessed by Grambsch-Therneau test for proportional hazards and evaluation for a non-zero slope of the scaled Schoenfeld residuals versus transformed time (**Supplementary Figure S6**). Based on inspection of the scaled Schoenfeld residuals, the model was stratified by tumor location. Influential observations were assessed with dfbeta and martingale residuals (**Supplementary Figure S7 and S8**). All tests were performed using R version 3.3.2 ^58^.

## DATA AVAILABILITY

Retrotransposon insertions and patient phenotypes are available in **Supplementary Table S1** and **S5**. WGS point mutations are available in EGA accession code EGAS00001003010.

## ACKNOWLEDGEMENTS

The authors thank Alison Ollikainen, Iina Vuoristo, Inga-Lill Åberg, Sini Marttinen, Marjo Rajalaakso, Sirpa Soisalo, Jiri Hamberg and Heikki Metsola for technical assistance and Alison Ollikainen also for proofreading the manuscript. This work was supported by grants from the Academy of Finland (Finnish Center of Excellence Program 2012-2017, 250345 and 2018-2025, 312041), The Finnish Cancer Society, The European Research Council (268648), The Sigrid Juselius Foundation; Jane and Aatos Erkko Foundation, SYSCOL (an EU FP7 Collaborative Project, 258236), the Nordic Information for Action eScience Center (NIASC) and Nordic Center of Excellence financed by NordForsk (Project number 62721). The following foundations are acknowledged for personal funding: Ida Montinin Säätio foundation, Cancer society of Finland, Juhani Ahon Foundation for Medical Research and The Maud Kuistila Memorial Foundation. The authors wish to acknowledge CSC-IT Center for Science, Finland, for computational resources.

## AUTHOR CONTRIBUTIONS

TC, PS and TT analyzed insertion data. TC, PS and EP contributed and performed insertion and transduction calling. TC, TT, UAH and JK prepared WGS samples. OK supervised WGS sample preparation. TC and OK contributed and organized RNA sample preparation. TC and TT performed statistical analysis. RK, EP, KP and NV were involved in primary WGS data analysis. TC and RK performed and analyzed insertion polyA/T calls and insertion orientation bias, and RK developed BasePlayer. AT and LF performed CIMP analysis. AT and KP designed and performed the analysis of insertion effect on gene expression. NV performed primary RNA seq analysis. AR and JB reviewed tumors. SK, AL, LRS, TK, TS and JPM provided patient samples. EK and MA contributed to the study design. TC, OK, EP, KP and LAA designed the study. OK, EP, KP and LAA supervised the study. All authors contributed to writing the manuscript.

